# TNFα reduces inhibitory transmission in young *Trem2^R47H^* Sporadic Alzheimer rats before observable Aβ and brain pathology

**DOI:** 10.1101/2020.08.20.256099

**Authors:** Siqiang Ren, Lionel Breuillaud, Wen Yao, Tao Yin, Kelly A. Norris, Simone P. Zehntner, Luciano D’Adamio

## Abstract

*Trem2^R47H^* rats, which carry the Alzheimer’s disease (AD) risk factor p.R47H variant of the microglia gene *TREM2* and produce human Aβ. Previously, we demonstrated that supraphysiological TNF-α boost glutamatergic transmission and suppresses Long-term-Potentiation (LTP), a surrogate of learning and memory, in peri-adolescent *Trem2^R47H^* rats (Ren et al., 2020). Here we tested the effect of the p.R47H *TREM2* variant on GABA transmission. We report that GABAergic transmission is decreased in *Trem2^R47H/R47H^* rats. This decrease is due to the acute and reversable action of TNF-α and is not associated whit changes in human Aβ levels and pathological brain lesions. Thus, the p.R47H *TREM2* variant changes the excitatory/inhibitory balance between glutamate and GABA transmission, favoring excitation. This unbalance could potentiate glutamate excitotoxicity and, over time, contribute to neuronal dysfunction, enhanced neuronal cells death and neurodegeneration. Future studies will determine whether this unbalance represents an early, Aβ-independent pathway leading to dementia.

## Introduction

Sporadic Alzheimer’s disease (SAD) represent ~95% of AD cases. Yet, the most commonly used animal organisms model Familial AD (FAD), which only represent ~5% of AD cases. This may be an issue because if FAD and SAD present significant pathogenic differences, therapeutic strategies effective in FAD animals may have limited therapeutic efficacy in SAD patients. Thus, model organisms that reproduce the pathogenesis of SAD would be helpful to identify therapeutic targets and test SAD-modifying therapeutics.

The p.R47H variant of the microglia gene *Triggering Receptor Expressed on Myeloid Cells 2* (*TREM2*) triples the risk of SAD in heterozygous carriers (Guerreiro, Wojtas et al. 2013). Like other SAD-associated *TREM2* variants, the p.R47H variant impairs the Aβ-clearing activities of microglia, presumably hampering elimination of toxic Aβ peptide forms (Zhou, Ulland et al. 2018, Zhou, Tan et al. 2019).

To study mechanisms by which this variant promotes neurodegeneration, we generated *Trem2^R47H^* knock-in (KI) rats. These rats carry the R47H mutation in the rat *Trem2* gene, exhibit normal *Trem2* splicing and expression and express two humanized *App* rat alleles that drive production of human Aβ (Tambini and D’Adamio 2020). Hence, this KI model is useful to study both human Aβ-dependent and Aβ-independent effects of the R47H mutation.

Pathogenic mechanisms leading to neurodegeneration and dementia may start early in life. To reveal early dysfunctions that may lead, over time, to neurodegeneration we have studied young *Trem2^R47H^* KI rats. Pre-adolescent (4 weeks old) and peri-adolescent (6-8 weeks old) *Trem2^R47H^* rats showed no alteration in steady-state levels of brain and cerebrospinal fluid (CSF) Aβ peptides (Ren, Yao et al. 2020, Tambini and D’Adamio 2020) suggesting that Aβ-clearance deficits caused by the p.R47H *TREM2* variant may manifest with aging. Yet, young *Trem2^R47H^* rats present significant increased brain and CSF concentrations of TNF-α, which cause augmented glutamatergic transmission as well as suppression of Long-term-Potentiation (LTP) (Ren, Yao et al. 2020), an electrophysiological surrogate of learning and memory. Physiological levels of TNF-α produced by microglia are necessary to maintain normal surface expression of AMPA receptors at post-synaptic termini; increased TNF-α concentrations promote rapid exocytosis of AMPA receptors in hippocampal pyramidal neurons increasing the strength of glutamatergic synaptic responses (Beattie, Stellwagen et al. 2002, Ogoshi, Yin et al. 2005, Stellwagen, Beattie et al. 2005, Stellwagen and Malenka 2006). The alterations in glutamatergic transmission found in *Trem2^R47H^* rats are consistent with these effects of TNF-α and establish a direct link between a pathogenic variant of the microglia specific *TREM2* gene and neuronal dysfunction of glutamatergic transmission and LTP.

In addition to boosting excitatory transmission, TNF-α decreases inhibitory synaptic strength by promoting endocytosis of γ-aminobutyric acid (GABA) receptors, hence reducing surface GABA receptors (Stellwagen, Beattie et al. 2005). Thus, in this study we tested whether the p.R47H *TREM2* variant may reduce GABA transmission *via* increased brain TNF-α levels with the purpose of determining whether the p.R47H *TREM2* variant changes the excitatory/inhibitory balance between glutamate and GABA transmission, favoring excitation. This unbalance could potentiate glutamate excitotoxicity and, over time, contribute to neuronal dysfunction, enhanced neuronal cells death and neurodegeneration.

## Results

### Reduced inhibitory synaptic transmission at hippocampal SC–CA3>CA1 synapses of peri-adolescent rats carrying the *Trem2*^*R47H*^ variant

We examined the effects of the *Trem2^R47H^* variant on GABAergic synaptic transmission in the hippocampal Schaeffer-collateral pathway. First, we examined paired-pulse facilitation (PPF) of GABA_A_ receptors postsynaptic current, which is inversely correlated to the presynaptic GABA release probability. Our results show that PPF with 50 and 200ms intervals is significantly increased in *Trem2^R47H/R47H^* rats (Figure 1A, B and C), indicating GABA release is undermined in *Trem2^R47H/R47H^* rats. Second, we analyzed miniature inhibitory postsynaptic currents (mIPSC). The frequency of mIPSCs that largely reflects presynaptic GAB_A_ release probability, is slightly decreased in both *Trem2^R47H/w^* and *Trem2^R47H/R47H^* rats; however, this decrease fails to achieve statistical significance (Figure 1D, E and H). Interestingly, the amplitude of mIPSCs that is dependent on levels of postsynaptic ionotropic GABA_A_ receptors was significantly decreased in both *Trem2^R47H/w^* and *Trem2^R47H/R47H^* rats (Figure 1D, F and I), suggesting a reduction of GABA_A_ receptors on the postsynaptic surface. Moreover, we found that in *Trem2^R47H/R47H^* rats, the decay time of mIPSC is shorter than in wild-type rats (Figure 1G). As the subunit composition of the GABA_A_ receptor determines the decay time of mIPSC (Eyre, Renzi et al. 2012), it is possible that the subunit composition of GABA_A_ receptor is altered in *Trem2^R47H/R47H^* rats. Altogether, these results suggest that the pathogenic variant p.R47H of the microglia gene *TREM2* leads to the reduction of GABAergic transmission to CA1 pyramidal neurons, and this effect is gene dosage dependent.

**Figure 1.**
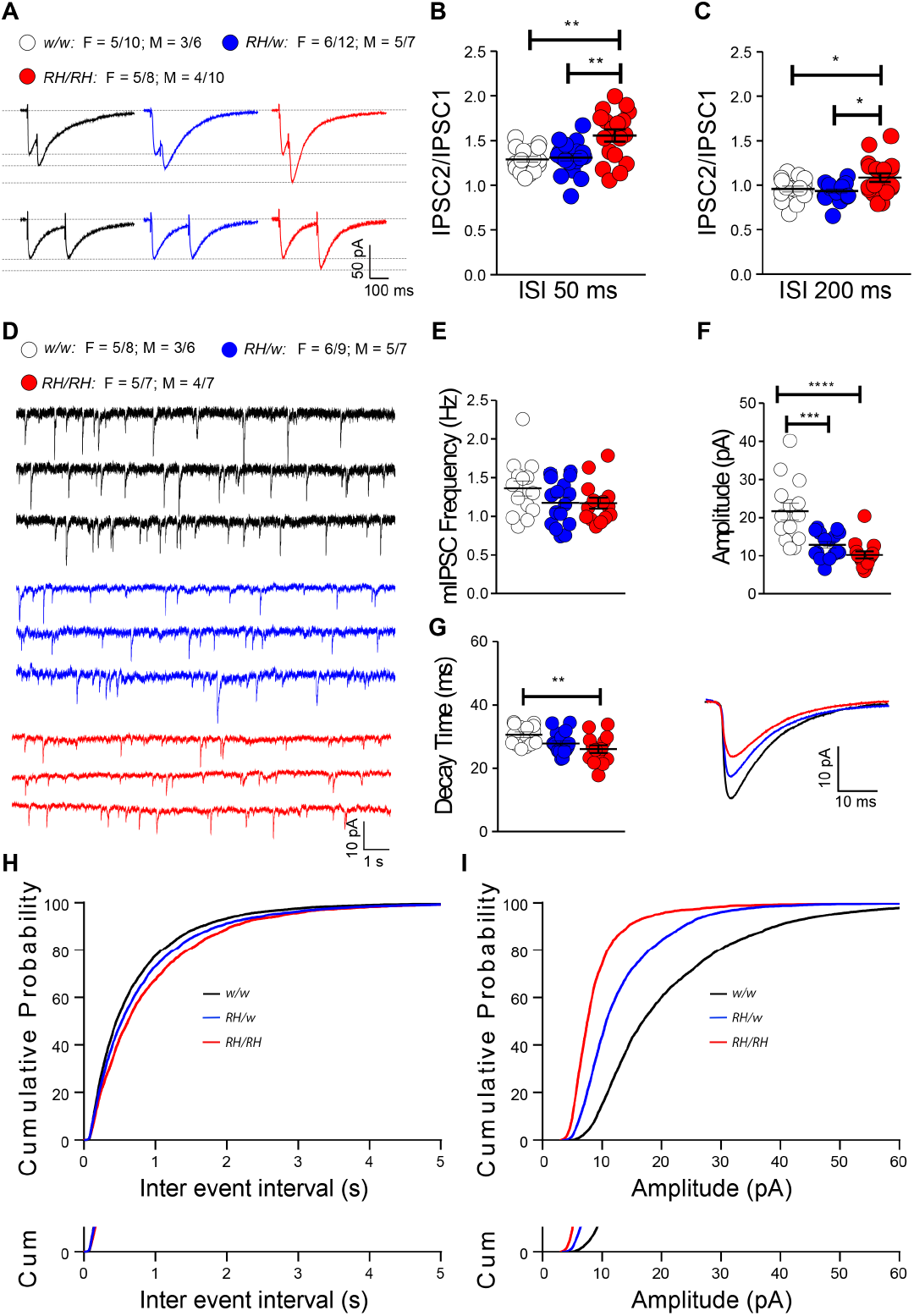
Inhibitory GABAergic synaptic transmission is decreased in *Trem2^R47H^* rats. **(A)** Representative traces of PPF of GABAergic transmission. **(B)** Plot of PPF at 50ms ISI and the representative traces. [F (2, 50)=8.968, P=0.0005****; post-hoc Tukey’s multiple comparisons test: *w/w vs. RH/w*, P=0.9485 (ns); *w/w vs. RH/RH*, P= 0.0014**; *RH/w vs. RH/RH*, P= 0.0022**]. **(C)** Plot of PPF at 200ms ISI. [F (2, 50) = 5.106, P=0.0096**; post-hoc Tukey’s multiple comparisons test: *w/w vs. RH/w*, P=0.9082 (ns); *w/w vs. RH/RH*, P= 0.0461*; *RH/w vs. RH/RH*, P=0.0118*]. **(D)** Representative traces of mini IPSCs. **(E)** Plot of the frequency of mini IPSCs. [F (2, 41) = 1.734, P=0.1893(ns)]. Notably, *RH* mutant rats show mIPSCs with decreased frequency, albeit this decrease did not reach a statistical significance. **(F)** Plot of the amplitude of mini IPSCs. [F (2, 41) = 17.04, P<0.0001****; post-hoc Tukey’s multiple comparisons test: *w/w vs. RH/w*, P= 0.0002 ***; *w/w vs. RH/RH*, P<0.0001****; *RH/w vs. RH/RH*, P= 0.3947(ns)]. **(G)** Plot of the decay time of mini IPSCs. [F (2, 41) = 5.254, P=0.0093**; post-hoc Tukey’s multiple comparisons test: *w/w vs. RH/w*, P= 0.1257(ns); *w/w vs. RH/RH*, P=0.0070**; *RH/w vs. RH/RH*, P= 0.3890 (ns)]. Representative averaged mini IPSCs traces are shown on right. Note that the amplitude and of decay time of mini IPSCs are significantly increased in RH mutant rats. **(H)** Plot of the cumulative probability of mini IPSCs inter-event intervals. **(I)** Plot of the cumulative probability of mini IPSCs amplitude. Data are represented as mean ± SEM and were analyzed by ordinary one-way ANOVA followed by post-hoc Tukey’s multiple comparisons test when ANOVA showed significant differences. For each type of recordings, we indicate the number of animals by genotype and sex, plus the number of recording by genotype and sex as follow: 1) genotypes: *w/w* = *Trem2^w/w^*, *RH/w* =*Trem2^R47H/w^*, *RH/RH* = *Trem2^R47H/R47H^;*2) sex: F = female, M = males; 3) number of animals and number of recordings from animals: n/n’, were n = number of animals, n’ = number of recordings from the n animals. For example, the *w/w*: F = 5/10; M = 3/6 in A indicates that data for PPF for the *Trem2^w/w^* rats were obtained from 5 females and 3 males, and that 10 recordings were obtained from the 5 females and 6 recordings from the 3 males.

### The reduced inhibitory synaptic transmission at hippocampal SC–CA3>CA1 synapses of peri-adolescent *Trem2^R47H^* rats is caused by supraphysiological TNF-α

TNF-α produced by glia is necessary for physiological post-synaptic surface expression of both AMPA and GABA receptors. These two opposite effects of physiological TNF-α cooperate in maintaining physiological excitatory/inhibitory balance and the excitatory synaptic strength. Increased TNF-α concentration causes a swift surface AMPA receptors expression at post-synaptic termini and endocytosis of GABA receptors in hippocampal pyramidal neurons, changing the excitatory/inhibitory balance and favoring excitation (Beattie, Stellwagen et al. 2002, Ogoshi, Yin et al. 2005, Stellwagen, Beattie et al. 2005, Stellwagen and Malenka 2006). Consistently, young *Trem2^R47H/R47H^* rats show increased levels of TNF-α in brain and CSF, which leads to enhanced glutamatergic transmission (Ren, Yao et al. 2020) To test whether TNF-α mediates the reduced inhibitory synaptic transmission at hippocampal SC–CA3>CA1 synapses of peri-adolescent rats carrying the *Trem2^R47H^* variant, we treated hippocampal slices with a neutralizing antibody to rat TNF-α (anti-TNF-α), which functions as a TNF-α antagonist. To control for off-target effects of the antibody, we used a Goat IgG isotype control. The 50% neutralization dose (ND_50_) of this anti-TNF-α antibody against the cytotoxic effect of recombinant rat TNF-α (0.25 ng/mL) is about 500 ng/ml. Since physiological levels of TNF-α are necessary for normal glutamatergic transmission and most of the activities of TNF-α can be rapidly reversed (Beattie, Stellwagen et al. 2002, Ogoshi, Yin et al. 2005, Stellwagen, Beattie et al. 2005, Stellwagen and Malenka 2006), we tested the acute effects 10ng/ml of anti-TNF-α, a concentration ~50 times lower than the ND_50_. At this concentration, anti-TNF-α occluded the increased PPF (Figure 2A, B and C), the decreased mIPSCs amplitude and decay time (Figure 2D, F, G and I) in *Trem2^R47H/R47H^* rats. The goat IgG isotype control did not restore inhibitory GABAergic transmission alterations observed in the mutant rats (Figure 2D, F, G and I) suggesting that the effects of anti-TNF-α are specific. In addition, these low doses of anti-TNF-α do not alter inhibitory transmission in *Trem2^w/w^* rats (Figure 2A-I), indicating that at least at this dosage, anti-TNF-α only targets GABAergic transmission alterations triggered by excess TNF-α set off by the *Trem2^R47H^* variant. Overall, these data indicate that the decrease of GABAergic transmission at SC–CA3>CA1 synapses of *Trem2^R47H/R47H^* rats is due to the acute action of supraphysiological TNF-α concentrations prompted by the *Trem2^R47H^* variant and are rapidly reversable.

**Figure 2.**
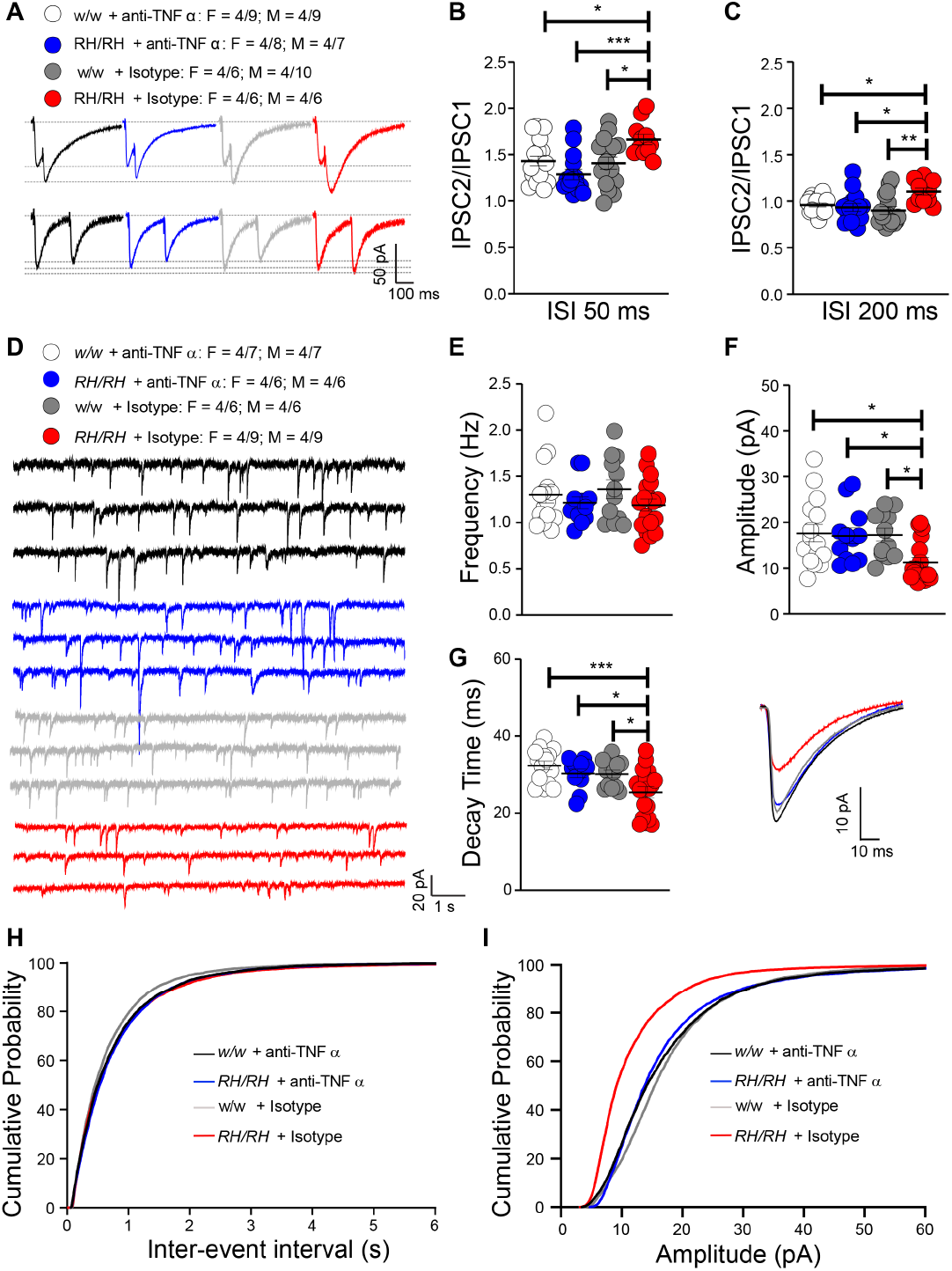
The reduced inhibitory GABAergic synaptic transmission in in *Trem2^R47H^* rats is restored by reducing TNFα function. **(A)** Representative traces of PPF of GABAergic transmission. Representative traces are averaged from 20 sweeps. **(B)** Plot of PPF at 50ms ISI. [F (3, 56) = 6.021, P=0.0013**; post-hoc Tukey’s multiple comparisons test: *w/w +* anti-TNF-α *vs. RH/RH* + anti-TNF-α, 0.2643 (ns); *w/w +* anti-TNF-α *vs. RH/RH* + Isotype, 0.0452*; *w/w +* anti-TNF-α *vs. w/w* + Isotype, 0.9897 (ns); *RH/RH +* anti-TNF-α *vs. RH/RH* + Isotype, P=0.0005***; *RH/RH +* anti-TNF-α *vs. w/w* + Isotype P=0.4452 (ns); *RH/RH +* Isotype *vs. w/w +* Isotype, P=0.0267*]. **(C)** Plot of PPF at 200ms ISI. [F (3, 56) = 5.361, P=0.0026**; post-hoc Tukey’s multiple comparisons test: *w/w +* anti-TNF-α *vs. RH/RH* + anti-TNF-α, 0.9485 (ns); *w/w +* anti-TNF-α *vs. RH/RH* + Isotype, 0.0348*; *w/w +* anti-TNF-α *vs. w/w* + Isotype, 0.5884 (ns); *RH/RH +* anti-TNF-α *vs. RH/RH* + Isotype, P=0.0128*; *RH/RH +* anti-TNF-α *vs. w/w* + Isotype P=0.9030 (ns); *RH/RH +* Isotype *vs. w/w +* Isotype, P=0.0017**]. Note that the increased in PPF of mini IPSCs at 50 and 200 ms are reversed by anti-TNFα antibody application in RH mutant rats. **(D)** Representative traces of mini IPSCs. **(E)** Plot of the frequency of mini IPSCs. [F (3, 53) = 0.9519, P=0.4223 (ns)]. **(F)** Plot of the amplitude of mini IPSCs. [F (3, 53) = 4.562, P=0.0065***; post-hoc Tukey’s multiple comparisons test: *w/w +* anti-TNF-α *vs. RH/RH* + anti-TNF-α, 0.9945 (ns); *w/w +* anti-TNF-α *vs. RH/RH* + Isotype, 0.0142*; *w/w +* anti-TNF-α *vs. w/w* + Isotype, 0.9989 (ns); *RH/RH +* anti-TNF-α *vs. RH/RH* + Isotype, P=0.0458*; *RH/RH +* anti-TNF-α *vs. w/w* + Isotype P=0.9996 (ns); *RH/RH +* Isotype *vs. w/w +* Isotype, P=0.0350*]. **(G)** Plot of the decay time of mini IPSCs. [F (3, 53) = 6.587, P=0.0007***; post-hoc Tukey’s multiple comparisons test: *w/w +* anti-TNF-α *vs. RH/RH* + anti-TNF-α, 0.6728 (ns); *w/w +* anti-TNF-α *vs. RH/RH* + Isotype, 0.0005***; *w/w +* anti-TNF-α *vs. w/w* + Isotype, 0.6144 (ns); *RH/RH +* anti-TNF-α *vs. RH/RH* + Isotype, P=0.0342*; *RH/RH +* anti-TNF-α *vs. w/w* + Isotype P=0.9997 (ns); *RH/RH +* Isotype *vs. w/w +* Isotype, P=0.0436*].The representative averaged traces of the mini IPSCs are shown on the right. Note that the decreased amplitude and decay time of mini IPSCs in RH /RH rats was restored by anti-TNFα antibody application. **(H)** Plot of the cumulative probability of mini IPSCs inter-event intervals. Plot of the cumulative probability of mini IPSCs amplitude. All data represent means ± SEM. Data were analyzed by Ordinary one-way ANOVA followed by post-hoc Tukey’s multiple comparisons test when ANOVA showed significant differences.

### Aβ peptides levels are not changed in the brain of peri-adolescent *Trem2^R47H^* rats

Analysis of brain homogenates from pre-adolescent rats (Tambini and D’Adamio 2020) showed no significant alterations in levels of human Aβ40, Aβ42 and the Aβ42/Aβ40 ratio in *Trem2^R47H^* rats, even though the Trem2^R47H^ variant reduces binding and clearance of human Aβ *in vitro* (Zhao, Wu et al. 2018). Previously, we found no changes in Aβ levels in the CSF of peri-adolescent animals as compared to *Trem2^w/w^* rats (Ren, Yao et al. 2020). However, CSF concentrations of Aβ may not reflect the brain Aβ levels since aggregation of Aβ peptides in brain parenchyma may influence Aβ levels in the CSF. Thus, we assessed further Aβ levels in the brains of *Trem2^R47H^* peri-adolescent animals, which were tested for glutamatergic (Ren, Yao et al. 2020) and GABAergic transmission (Figures 1 and 2). No differences were seen in Aβ38, Aβ40, Aβ42 levels and the Aβ42/Aβ40 ratio between peri-adolescent *Trem2^w/w^*, *Trem2^R47H/w^*, and *Trem2^R47H/R47H^* rats (Figure 3), further suggesting that the reduced Aβ clearance caused by the Trem2^R47H^ variant *in vitro* does not result in significant alterations of Aβ steady-state levels *in vivo*, at least in pre- and peri-adolescent rats.

**Figure 3.**
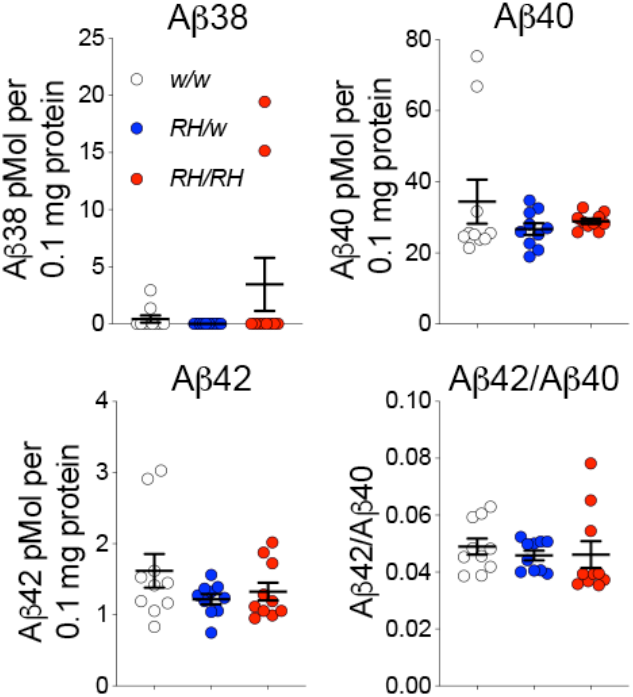
Levels of human Aβ species are similar in the brain of peri-adolescent *Trem2^w/w^*, *Trem2^R47H/w^*, and *Trem2^R47H/R47H^* rats. Levels of Aβ38, Aβ40, and Aβ42/Aβ40 ratio in 7-8 weeks old *Trem2^w/w^*, *Trem2^R47H/w^*, and *Trem2^R47H/R47H^* rat brains. We used 5 males and 5 females for each genotype. Data are represented as mean ± SEM. Data were analyzed by ordinary one-way ANOVA. No differences were seen in Aβ38 [F_(2, 27)_ = 1.931, P=0.1645], Aβ40 [F_(2, 27)_ = 1.132, P=0.3374], Aβ42 [F_(2, 27)_ = 1.668, P=0.2074] levels and the Aβ42/Aβ40 ratio [F_(2, 27)_ = 0.2683, P=0.7667].

### *Trem2^R47H/w^* and *Trem2^R47H/R47H^* adult (3 months old) rat brains show no evidence of Aβ aggregation, neurodevelopmental or histopathological changes

The Aβ ELISA may not efficiently measure aggregated insoluble Aβ species. These species may trigger TNF-α release and impact neurodevelopment and/or cause overt neuropathology. To test these possibilities, we used histology and immunohistochemistry (IHC) to characterize brains from 3-month-old male and female *Trem2^w/w^*, *Trem2^R47H/w^* and *Trem2^R47H/R47H^* rats (see **Table 1**). We tested 3 months-old rats to increase the possibility of detecting pathology that may start in pre-adolescent rats but may be detectable by histology and IHC only weeks later. Regions of analysis included the frontal cortex, cingulate cortex, whole hippocampus, and entorhinal cortex. No gross morphological changes were evident by hematoxylin and eosin (H&E) staining in any of the rats analyzed (Figure 4). Qualitative inspection of NeuN staining showed no appreciable changes in neuronal density in any of the regions analyzed. Qualitative analysis performed on the hippocampus (CA1) and the somatosensory cortex (Cx) did not indicate overt neuronal loss in *Trem2^R47H/w^* and *Trem2^R47H/R47H^* rats as compared to *Trem2^w/w^* rats (Figure 5). Despite the presence of elevated proinflammatory cytokines in the brain and CSF of *Trem2^R47H/R47H^* and to a lesser extent of *Trem2^R47H/w^* rats, no evidence of significant astrocytosis or microgliosis was observed (Figure 4, 5 and 6). As evaluated by the staining intensity (Figure 5) and cellular morphology of GFAP and IBA1 stained tissues (Figure 6 A&B). While a trend toward higher IBA-1 staining intensity was observed in the Trem2^R47H/R47H^ rats as compared to Trem2^w/w^ and *Trem2^R47H/w^* rats, this was not statistically significant. The microglia presented numerous fine processes, characteristic of the resting state, and did not present obvious intermediate or amoeboid morphologies with enlarged processes or cell bodies (Figure 6A) in all three genotypes. Similarly, astrocytes did not present hypertrophy of soma and processes in any of the groups (Figure 6B). Amyloid plaques, as measured by simultaneous co-staining with the anti-Aβ antibodies 6E10 and 4G8 were absent in all tissue analyzed (Figure 4). Those results are consistent with the similar Aβ40 or Aβ42 levels in *Trem2^w/w^*, *Trem2^R47H/w^* and *Trem2^R47H/R47H^* rats as well as the absence of plaque in 3 month rats with humanized Aβ (Tambini, Yao et al. 2019). Moreover, Tau phosphorylation as measured by AT8 immunostaining was absent in all the groups (Figure 4), and a modified Bielschowsky silver staining did not reveal plaques, dystrophic neurites or axonal pathologies in any of the tissue analyzed (Figure 4). Overall, histological analysis of these rats shows no obvious evidence of neurodevelopmental impairments, neurodegeneration, neuroinflammation, or AD-like pathology at 3 months of age.

**Figure 4.**
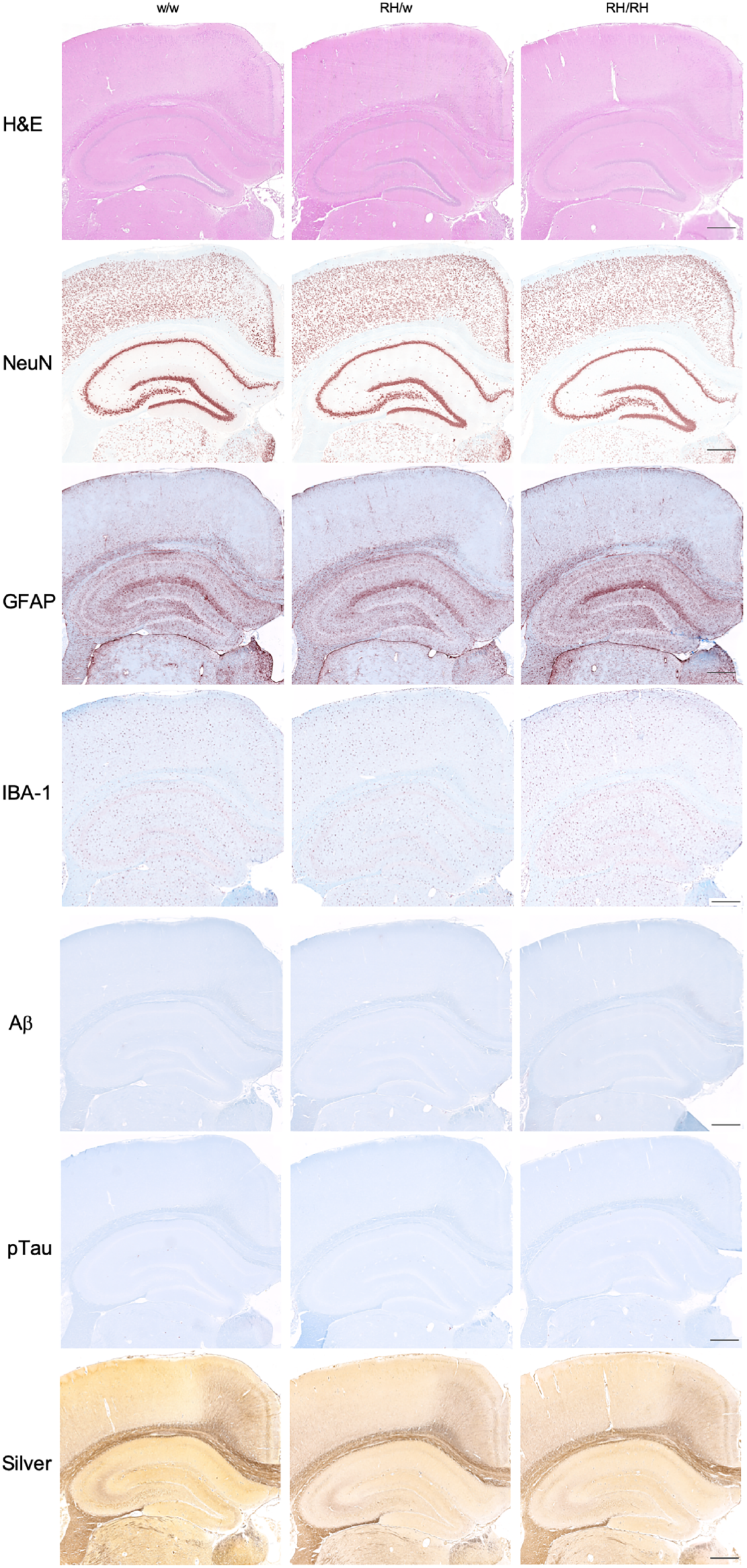
Immunohistochemistry staining in 3-month-old *Trem2^w/w^*, *Trem2^R47H/w^*, and *Trem2^R47H/R47H^* rats. Representative images of the anterior hippocampus and overlaying somatosensory cortex of 3-month-old male *Trem2^w/w^*, *Trem2^R47H/w^*, and *Trem2^R47H/R47H^* rat brains. Illustrates, from top to bottom, H&E, NeuN, GFAP, IBA-1, Aβ, pTau and Bielschowski Silver staining respectively. No observable differences morphology (H&E) or neuronal (NeuN) cellularity are observed. The staining intensity of the microglial (IBA-1) and astrocytic (GFAP) markers are similar across all three genotypes. While no Aβ or pTau expression can be observed and no Bielschowski Silver stained plaques or tangles are present. Immunohistochemistry staining were performed on: *Trem2^w/w^* (4 males and 5 females), *Trem2^R47H/w^* (4 males and 4 females) and *Trem2^R47H/R47H^* (4 males and4 females). The scale bar is equivalent to 500 microns.

**Figure 5.**
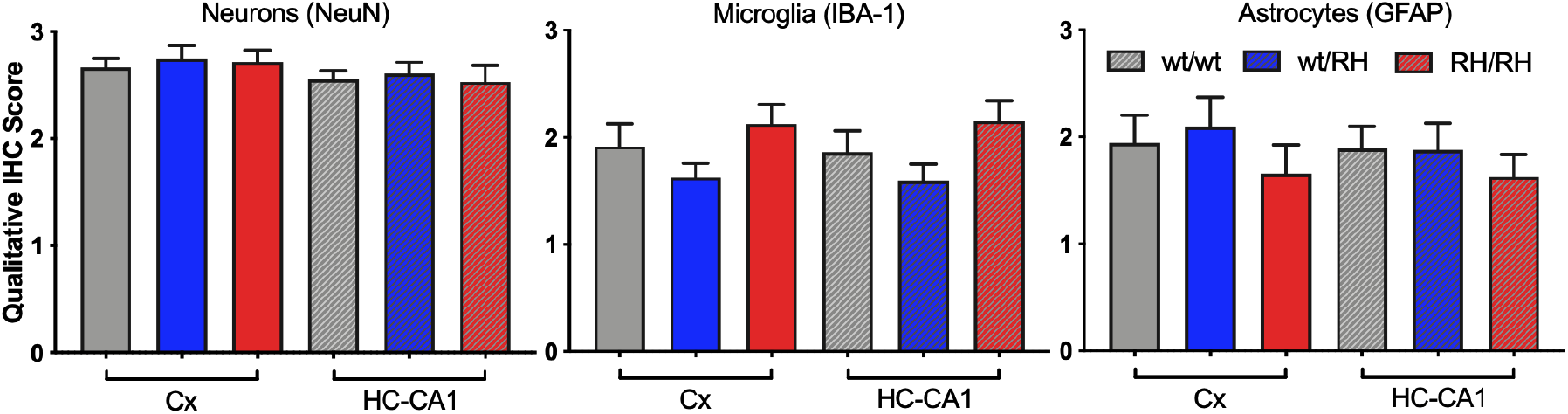
Qualitative assessment of the NeuN, IBA-1 and GFAP staining in 3-month-old *Trem2^w/w^*, *Trem2^R47H/w^*, and *Trem2^R47H/R47H^* rats. Immunohistochemistry staining was score in: *Trem2^w/w^* (4 males and 5 females), *Trem2^R47H/w^* (4 males and 4 females) and *Trem2^R47H/R47H^* (4 males and 4 females). Data are represented as mean ± SEM of the qualitative score within the cortex and hippocampus-CA1 regions. Data were analyzed by ordinary one - way ANOVA within each brain region. No statistically significant differences were seen in NeuN_Cx_ [F_(2,21)_=1.736, P=0.200], NeuN_HC-CA1_ [F_(2,21)_= 0.533,P=0.594], IBA-1_Cx_ [F_(2,22)_=0.375, P=0.692], IBA-1_HC-CA1_ [F_(2,22)_= 0.507, P=0.609], GFAP_Cx_ [F_(2,22)_=0.159, P= 0.854], or GFAP_HC-CA1_ [F_(2,22)_=0.0237,P=0.977].

**Figure 6.**
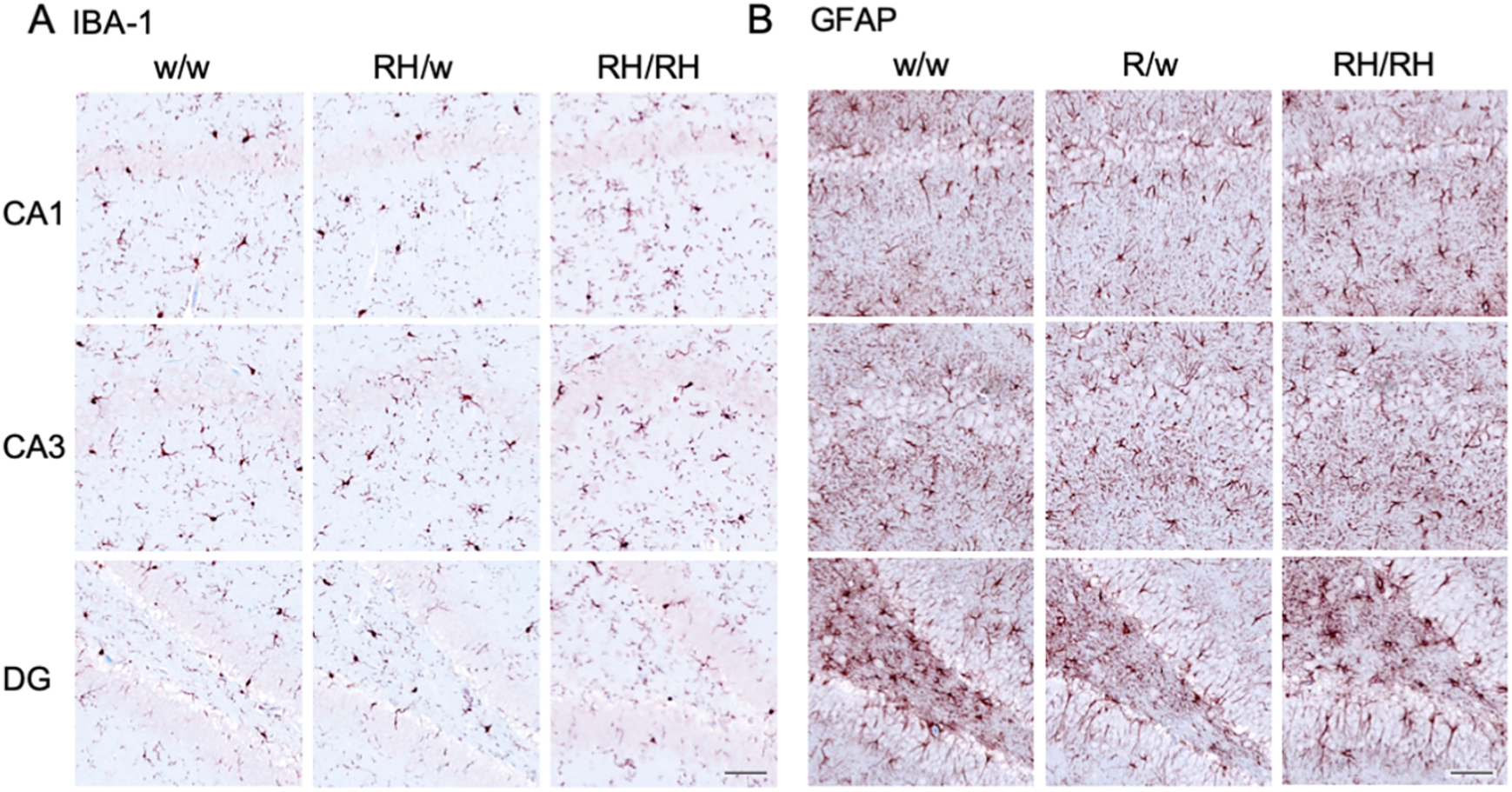
Astrocyte and microglia Immunohistochemistry staining in 3 month old *Trem2^w/w^*, *Trem2^R47H/w^*, and *Trem2^R47H/R47H^* rats. Representative images illustrate microglia (**A, IBA-1**) and astrocytes (**B, GFAP**) staining with a red-brown chromagen in the dorsal hippocampus-CA1, hippocampus-CA3, and hippocampus-dentate gyrus (DG), of 3-month-old male *Trem2^w/w^*, *Trem2^R47H/w^*, and *Trem2^R47H/R47H^* rat brains (left to right). The scale bar is equivalent to 500 microns.

## DISCUSSION

Pro-inflammatory cytokines, especially TNF-α, are significantly increased in the brain and CSF of young *Trem2^R47H^* rats (Ren, Yao et al. 2020). Consistent with the evidence that: 1) TNF-α produced by glia controls post-synaptic physiological expression of AMPA receptors; 2) increased concentrations of TNF-α cause rapid AMPA receptors exocytosis increasing excitatory synaptic strength (Beattie, Stellwagen et al. 2002, Ogoshi, Yin et al. 2005, Stellwagen, Beattie et al. 2005, Stellwagen and Malenka 2006); we found that supraphysiological TNF-α concentrations boost glutamatergic transmission and suppress LTP, a surrogate of learning and memory, in peri-adolescent *Trem2^R47H^* rats (Ren, Yao et al. 2020).

TNF-α also physiologically regulates, in an opposite manner, inhibitory synaptic strength by promoting endocytosis GABA receptors, hence reducing surface GABA receptors at inhibitory synapses (Stellwagen, Beattie et al. 2005). In accord, we show here that young *Trem2^R47H^* rats have reduced GABA responses (Figure 1). Low doses of a neutralizing anti-TNF-α antibody occludes these alterations (Figure 2) indicating that supraphysiological TNF-α concentrations impair GABAergic transmission in *Trem2^R47H^* rats. This evidence also indicates that GABAergic deficits must be due to an acute and constant action of supraphysiological TNF-α.

TNF-α-dependent synaptic transmission alterations occur in the absence of changes in steady-state levels of soluble Aβ (Figure 3 and (Ren, Yao et al. 2020, Tambini and D’Adamio 2020) and obvious evidence of neurodevelopmental impairments, neurodegeneration, neuroinflammation, or AD-like pathology (Figures 4-6). Overall, these data suggest that the TNF-α-dependent synaptic transmission alterations caused by the p.R47H *TREM2* variant are independent of, and perhaps precede, changes in Aβ steady-state levels and brain pathology.

Since in the brain *TREM2* expression is restricted to microglia (Schmid, Sautkulis et al. 2002), it is plausible that *Trem2^R47H^*-microglia may be the source of supraphysiological TNF-α. *Trem2^R47H^*-microglia may also promote TNF-α production by other cell types -such as astrocytes. The possibilities that non-brain resident cells may be the source of extra TNF-α or that TNF-α clearance is altered in *Trem2^R47H^* rats, cannot be discounted. These possibilities do not need to be mutually exclusive. Interestingly, while these observations may suggest that increased levels of TNF-α and other cytokines measured in *Trem2^R47H/R47H^* is attributable to microglia or astrocytes, their immunoactivity does not appear to be associated with obvious morphological changes. However, we cannot exclude a mildly activated microglia phenotype, characterized by increased branching of thin processes as well as lengthening of processes and the secretion of proinflammatory cytokines. Studies using a model of prion disease (Perry, Cunningham et al. 2007) have indicated that microglia can switch to a phenotype contributing to neuronal damage without morphological changes (Perry, Cunningham et al. 2007). Thus, in some experimental models of CNS disease there is no direct correlation between morphological profile and functional phenotype in microglia.

SAD is a disease of old age. Therefore it is reasonable to assume that changes in the excitatory/inhibitory balance between glutamate (Ren, Yao et al. 2020) and GABA transmission (this paper), favoring excitation, caused by the p.R47H *TREM2* variant have no relevance to SAD pathogenesis. However, it is possible that these early dysfunctions may potentiate glutamate excitotoxicity enhancing neuronal cells death and culminate into obvert cognitive deficits, brain pathology and neurodegeneration decades later. In this context, it is worth mentioning that several genes linked to dementia, including *APP*, *PSEN1*, *PSEN2* and *ITM2b* play a physiological role in glutamatergic transmission, and that mutations linked to familial dementia alter this physiological functions (Fotinopoulou, Tsachaki et al. 2005, Matsuda, Giliberto et al. 2005, Matsuda, Giliberto et al. 2008, Norstrom, Zhang et al. 2010, Tamayev, Giliberto et al. 2010, Tamayev, Matsuda et al. 2010, Groemer, Thiel et al. 2011, Matsuda, Matsuda et al. 2011, Kohli, Pflieger et al. 2012, Wu, Yamaguchi et al. 2013, Del Prete, Lombino et al. 2014, Fanutza, Del Prete et al. 2015, Lundgren, Ahmed et al. 2015, Xia, Watanabe et al. 2015, Tambini, Yao et al. 2019, Yao, Tambini et al. 2019, Yao, Yin et al. 2019).

In conclusion, our studies indicate that the pathogenic p.R47H *TREM2* variant changes the excitatory/inhibitory balance favoring excitation, that these changes happen early in life, are dependent on supraphysiological TNF-α and independent of changes in Aβ steady-state levels and brain pathology (Figure 7). More studies will be needed to determine whether this microglia-neuronal axis contributes to SAD pathogenesis in humans.

**Figure 7.**
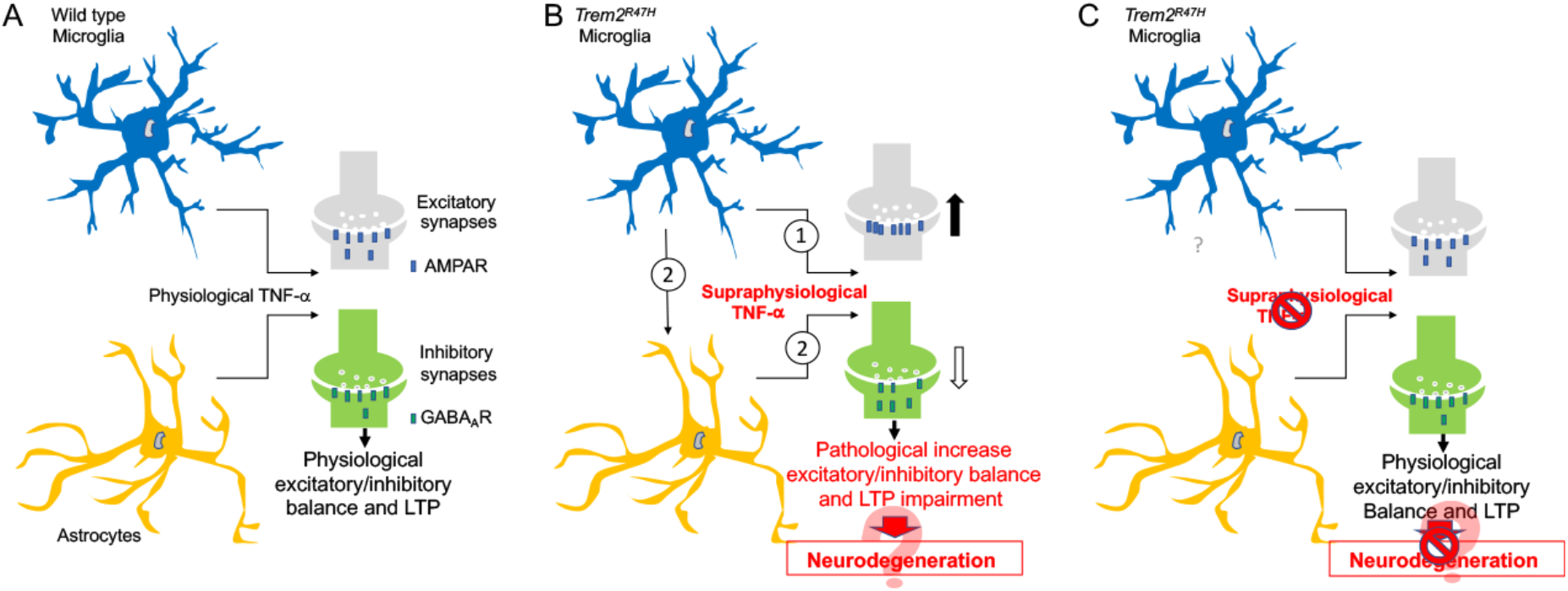
Model depicting how the p.R47H TREM2 variant may enhance the excitatory/inhibitory balance. **(A)** Astrocytes and microglia set physiological excitatory/inhibitory balance and LTP *via* TNF-α. (**B**) *TREM2* expression is restricted to microglia: thus, it is likely that microglia expressing the p.R47H variant are the source of supraphysiological TNF-α①. *Trem2^R47H^*-microglia may also promote TNF-α production by other cell types, such as astrocytes②. These two possibilities are not mutually exclusive. Supraphysiological TNF-α impairs LTP and increases the excitatory/inhibitory balance by enhancing glutamate transmission and reducing GABA transmission. A swift surface exocytosis of AMPA receptors and endocytosis of GABA receptors at post-synaptic termini could be one mechanism by which supraphysiological TNF-α increases the excitatory/inhibitory balance and reduces LTP. These early dysfunctions may enhance glutamate excitotoxicity and neuronal cells death, culminating into obvert cognitive deficits, brain pathology and neurodegeneration decades later. (**C**) Resetting TNF-α activity at physiological levels normalizes excitatory/inhibitory balance and LTP, and could prevent neurodegeneration.

## Materials and Methods

| Key Resources Table |  |  |  |  |
| --- | --- | --- | --- | --- |
| Reagent type (species) or resource | Designation | Source or reference | Identifiers | Additional information |
| genetic reagent (Rattus Norvegicus) | *App<sup>h</sup>* | (Tambini, Yao et al. 2019) | | Rat App allele with humanize Aβ region |
| genetic reagent (Rattus Norvegicus) | *Trem2<sup>R47H</sup>* | (Tambini and D'Adamio 2020) |  | Rat Trem2 allele with R47H mutation |
| commercial assay or kit | V-PLEX Plus Aβ Peptide Panel 1 | Meso Scale Discovery | Cat# K15200G | Used following manufacturer's recommendations |
| antibody | Polyclonal Goat IgG anti-Rat TNFα | R and D Systems | RRID:AB_354511 | 10 ng/ml in ACSF |
| antibody | Polyclonal Goat IgG. antibody | R and D Systems | RRID:AB_354267 | 10 ng/ml in ACSF |
| antibody | Anti-NeuN Mouse monoclonal A60 | Millipore | RRID:AB_431321 | Targets neurons: 1:3000 dilution. Antigen retrieval: Citrate pH 6.0 HIER |
| antibody | Anti-Aβ Mouse monoclonal 6E10 | Biolegend | RRID:AB_2565327 | Targets Aβ amino acids 1-16: 1:1000 dilution. Antigen retrieval: 80% Formic acid |
| antibody | Anti-Aβ Mouse monoclonal 4G8 | Biolegend | RRID:AB_2734547 | Targets Aβ amino acids 17-24: 1:1000 dilution. Antigen retrieval: 80% Formic acid |
| antibody | Anti-IBA-1, Rabbit polyclonal | Wako | RRID:AB_839504 | Targets Microglia: 1:2000 dilution. Antigen retrieval: Citrate pH 6.0 HIER |
| antibody | Anti-GFAP, Rabbit polyclonal Antibody | Thermo Fisher Scientific | RRID:AB_2109797 | Targets Astrocytes: 1:2000 dilution. Antigen retrieval: Citrate pH 6.0 HIER |
| antibody | Anti-Phospho-Tau (Ser202, Thr205), Mouse monoclonal | Thermo Fisher Scientific | RRID:AB_223647 | Targets NFTs: 1:2000 dilution. Antigen retrieval: Citrate pH 6.0 HIER |
| antibody | Rabbit anti-Mouse | Jackson Immuno research Labs | RRID:AB_2340038 | IHC secondary amplification reagent used at 1:200 |
| antibody | Donkey anti-Rabbit-bio | Jackson Immuno research Labs | RRID:AB_2340593 | Biotinylated IHC secondary amplification reagent used at 1:200 |
| antibody | Goat anti-Rabbit-HRP | Jackson Immuno research Labs | RRID:AB_2307391 | Peroxidase conjugated IHC secondary amplification reagent used at 1:200 |
| antibody | Streptavidin-HRP | Abcam | Cat#ab64269 | SA conjugated IHC reagent (ready to use) |
| software | LinRegPCR s | hartfaalcentrum.nl |  |  |
| software | pCLAMP10 | Molecular Devices |  |  |
| software | Image Lab | Biorad | RRID:SCR_014210 |  |
| software | GraphPad Prism |  | RRID:SCR_002798 |  |

### Rats and ethics statement

All experiments were done according to policies on the care and use of laboratory animals of the Ethical Guidelines for Treatment of Laboratory Animals of the NIH. The procedures were described and approved by the Rutgers Institutional Animal Care and Use Committee (IACUC) (protocol number 201702513). All efforts were made to minimize animal suffering and reduce the number of animals used. The animals were housed two per cage under controlled laboratory conditions with a 12 hr dark light cycle, a temperature of 22 ± 2°C. Rats had free access to standard rodent diet and tap water.

### Brain slice preparation

6-8-week rats were deeply anesthetized with isoflurane and then intracardially perfused with 20 ml ice-cold cutting solution containing (in mM) 120 Choline Chloride, 2.6 KCl, 26 NaHCO_3_, 1.25 NaH_2_PO_4_, 0.5 CaCl_2_, 7 MgCl_2_, 1.3 Ascorbic Acid, 15 Glucose. The brains were rapidly removed from the skull. Coronal brain slices containing the hippocampal formation (400 μm thick) were prepared in the ice-cold cutting solution bubbled with 95% O_2_/5% CO_2_ using Vibratome VT1200S (Leica Microsystems, Germany). The slices were transferred into an interface chamber in ACSF containing (in mM): 126 NaCl, 3 KCl, 1.2 NaH_2_PO_4_; 1.3 MgCl_2_, 2.4 CaCl_2_, 26 NaHCO_3_, and 10 glucose (at pH 7.3), bubbled with 95% O_2_ and 5% CO_2_, and incubated at 30 °C for 1 hour and then kept at room temperature afterwards. The hemi-slices were transferred to a recording chamber perfused with ACSF at a flow rate of ~2 ml/min using a peristaltic pump. Experiments were performed at 28.0 ± 0.1°C.

### Electrophysiological recording

Whole-cell recordings in the voltage-clamp mode (−70 mV) were made with patch pipettes containing (in mM): 135 KCl, 2 MgCl_2_, 0.1 EGTA, 10 HEPES, 2 Na_2_ATP, 0.2 Na_2_GTP and 5 QX-314 (PH 7.3, osmolarity 290–310 mOsm). Patch pipettes (resistance, 8–10 MΩ) were pulled from 1.5 mm thin-walled borosilicate glass (Sutter Instruments, Novato, CA) on a horizontal puller (model P-97; Sutter Instruments, Novato, CA).

CA1 neurons were viewed under upright microscopy (FN-1, Nikon Instruments, Melville, NY) and recorded with Axopatch-700B amplifier (Molecular Devices, San Jose, CA). Data were low-pass filtered at 2 kHz and acquired at 5–10 kHz. The series resistance (Rs) was consistently monitored during recording in case of reseal of ruptured membrane. Cells with Rs >20 MΩ or with Rs deviated by >20% from initial values were excluded from analysis. Basal postsynaptic synaptic responses were evoked at 0.05 Hz by electrically stimulating the Schaffer collateral afferents using concentric bipolar electrodes. Inhibitory postsynaptic currents (IPSCs) were recorded with membrane potential held at −70 mV in ACSF containing 10 μM NBQX to block AMPA receptor current. The stimulation intensity was adjusted to evoke IPSCs that were 40% of the maximal evoked amplitudes (“test intensity”). For recording of paired-pulse ratio (PPR), paired-pulse stimuli with 50ms or 200ms inter-pulse interval were given at test intensity. The PPR was calculated as the ratio of the second IPSC amplitude to the first. Mini IPSCs were recorded by maintaining neurons at −70 mV with ACSF containing 1μM TTX to block action potentials, and μM NBQX to block AMPA receptor current. Mini IPSCs were recorded for 5-10 mins for analysis. Data were collected and analyzed using the Axopatch 700B amplifiers and pCLAMP10 software (Molecular Devices) and mEPSCs are analyzed using mini Analysis Program.

### Antibodies treatment

10ng/ml Goat anti-TNF-α (AF-510-NA, R&D Systems) or Goat IgG control (AB-108-C, R&D Systems) was incubated right after slice cutting and perfused throughout recordings. Experiments were performed at 28.0 ± 0.1°C.

### Rat brain preparation for ELISA

Rats were anesthetized with isoflurane and perfused via intracardiac catheterization with ice-cold PBS. This perfusion step eliminates cytokines and Aβ present in blood. Brains were extracted and homogenized using a glass-teflon homogenizer (w/v =100 mg tissue/1 ml buffer) in 250 mM Sucrose, 20 mM Tris-base pH 7.4, 1 mM EDTA, 1mM EGTA plus protease and phosphatase inhibitors (ThermoScientific), with all steps carried out on ice or at 4°C. Total lysate was solubilized with 0.1% SDS and 1% NP-40 for 30 min rotating. Solubilized lysate was spun at 20,000 g for 10 min, the supernatant was collected and analyzed by ELISA.

### ELISA

Aβ38, Aβ40, and Aβ42 were measured with V-PLEX Plus Aβ Peptide Panel 1 6E10 (K15200G) and V-PLEX Plus Aβ Peptide Panel 1. Measurements were performed according to the manufacturer’s recommendations. Plates were read on a MESO QuickPlex SQ 120. Data were analyzed using Prism software and represented as mean ± SEM.

#### Tissue Preparation and Staining

Rat brain tissue was fixed and stored in 70% ethanol following transcardiac perfusion with PBS and 4% paraformaldehyde fixative. All tissues were dehydrated through graded ethanol and xylene, infiltrated with paraffin wax, and embedded in paraffin blocks. Slides were manually de-paraffinized and re-hydrated prior to the automated immunohistochemistry. Slides initially underwent antigen retrieval, by one of the following methods, heat-induced epitope-retrieval (HIER), HIER and Proteinase K (PK) treatment, or Formic acid treatment. HIER was performed by incubation in a citrate buffer (pH 6.0) and heating to 100^°^C for a period of 60 minutes. When performed before the 10min PK treatment, citrate HIER was limited to 20min. Formic acid treatment was a 10 min incubation in 80% formic acid, followed by washing in TBS-T. All IHC studies were performed at room temperature on a Lab Vision Autostainer. Briefly, slides were incubated sequentially with hydrogen peroxide for 5 minutes, to quench endogenous peroxidase, followed by 5 minutes in Protein Block, and then incubated with primary antibodies as outlined in the **Key Resources Table**. Antibody binding was amplified using the appropriate secondary reagents (20 minutes), followed by a HRP-conjugate (20 minutes), and visualized using the AEC chromogen (20 minutes). All IHC sections were counterstained with Acid Blue 129 and mounted with aqueous mounting medium (Zehntner, Chakravarty et al. 2008).

### Statistical analysis

Data were analyzed using GraphPad Prism software and expressed as mean ± s.e.m. Statistical tests used to evaluate significance are shown in Figure legends. Significant differences were accepted at P<0.05.

## REFERENCES

Beattie, E. C., et al. (2002). “Control of synaptic strength by glial TNFalpha.” Science 295(5563): 2282–2285.

Del Prete, D., et al. (2014). “APP is cleaved by Bace1 in pre-synaptic vesicles and establishes a pre-synaptic interactome, via its intracellular domain, with molecular complexes that regulate pre-synaptic vesicles functions.” PLoS One 9(9): e108576.

Eyre, M. D., et al. (2012). “Setting the time course of inhibitory synaptic currents by mixing multiple GABA(A) receptor α subunit isoforms.” J Neurosci 32(17): 5853–5867.

Fanutza, T., et al. (2015). “APP and APLP2 interact with the synaptic release machinery and facilitate transmitter release at hippocampal synapses.” Elife 4: e09743.

Fotinopoulou, A., et al. (2005). “BRI2 interacts with amyloid precursor protein (APP) and regulates amyloid beta (Abeta) production.” J Biol Chem 280(35): 30768–30772.

Groemer, T. W., et al. (2011). “Amyloid precursor protein is trafficked and secreted via synaptic vesicles.” PLoS One 6(4): e18754.

Guerreiro, R., et al. (2013). “TREM2 variants in Alzheimer’s disease.” N Engl J Med 368(2): 117–127.

Kohli, B. M., et al. (2012). “Interactome of the amyloid precursor protein APP in brain reveals a protein network involved in synaptic vesicle turnover and a close association with Synaptotagmin-1.” J Proteome Res 11(8): 4075–4090.

Lundgren, J. L., et al. (2015). “ADAM10 and BACE1 are localized to synaptic vesicles.” J Neurochem.

Matsuda, S., et al. (2005). “The familial dementia BRI2 gene binds the Alzheimer gene amyloid-beta precursor protein and inhibits amyloid-beta production.” J Biol Chem 280(32): 28912–28916.

Matsuda, S., et al. (2008). “BRI2 inhibits amyloid beta-peptide precursor protein processing by interfering with the docking of secretases to the substrate.” J Neurosci 28(35): 8668–8676.

Matsuda, S., et al. (2011). “Maturation of BRI2 generates a specific inhibitor that reduces APP processing at the plasma membrane and in endocytic vesicles.” Neurobiol Aging 32(8): 1400–1408.

Norstrom, E. M., et al. (2010). “Identification of NEEP21 as a ss-amyloid precursor protein-interacting protein in vivo that modulates amyloidogenic processing in vitro.” J Neurosci 30(46): 15677–15685.

Ogoshi, F., et al. (2005). “Tumor necrosis-factor-alpha (TNF-alpha) induces rapid insertion of Ca2+-permeable alpha-amino-3-hydroxyl-5-methyl-4-isoxazole-propionate (AMPA)/kainate (Ca-A/K) channels in a subset of hippocampal pyramidal neurons.” Exp Neurol 193(2): 384–393.

Perry, V. H., et al. (2007). “Systemic infections and inflammation affect chronic neurodegeneration.” Nat Rev Immunol 7(2): 161–167.

Ren, S., et al. (2020). “Microglia *TREM2^R47H^* Alzheimer-linked variant enhances excitatory transmission and reduces LTP via increased TNF-α levels.” Elife 9.

Schmid, C. D., et al. (2002). “Heterogeneous expression of the triggering receptor expressed on myeloid cells-2 on adult murine microglia.” J Neurochem 83(6): 1309–1320.

Stellwagen, D., et al. (2005). “Differential regulation of AMPA receptor and GABA receptor trafficking by tumor necrosis factor-alpha.” J Neurosci 25(12): 3219–3228.

Stellwagen, D. and R. C. Malenka (2006). “Synaptic scaling mediated by glial TNF-alpha.” Nature 440(7087): 1054–1059.

Tamayev, R., et al. (2010). “Memory deficits due to familial British dementia BRI2 mutation are caused by loss of BRI2 function rather than amyloidosis.” J Neurosci 30(44): 14915–14924.

Tamayev, R., et al. (2010). “Danish dementia mice suggest that loss of function and not the amyloid cascade causes synaptic plasticity and memory deficits.” Proc Natl Acad Sci U S A 107(48): 20822–20827.

Tambini, M. D. and L. D’Adamio (2020). “Trem2 Splicing and Expression are Preserved in a Human Abeta-producing, Rat Knock-in Model of Trem2-R47H Alzheimer’s Risk Variant.” Sci Rep 10(1): 4122.

Tambini, M. D. and L. D’Adamio (2020). “Trem2 Splicing and Expression are Preserved in a Human Aβ-producing, Rat Knock-in Model of Trem2-R47H Alzheimer’s Risk Variant.” Sci Rep 10(1): 4122.

Tambini, M. D., et al. (2019). “Facilitation of glutamate, but not GABA, release in Familial Alzheimer’s APP mutant Knock-in rats with increased beta-cleavage of APP.” Aging Cell 18(6): e13033.

Wu, B., et al. (2013). “Presenilins regulate calcium homeostasis and presynaptic function via ryanodine receptors in hippocampal neurons.” Proc Natl Acad Sci U S A 110(37): 15091–15096.

Xia, D., et al. (2015). “Presenilin-1 Knockin Mice Reveal Loss-of-Function Mechanism for Familial Alzheimer’s Disease.” Neuron 85(5): 967–981.

Yao, W., et al. (2019). “Tuning of glutamate, but not GABA, release by an intra-synaptic vesicles APP domain whose function can be modulated by beta-or alpha-secretase cleavage.” J Neurosci.

Yao, W., et al. (2019). “The Familial dementia gene ITM2b/BRI2 facilitates glutamate transmission via both presynaptic and postsynaptic mechanisms.” Sci Rep 9(1): 4862.

Zehntner, S. P., et al. (2008). “Synergistic tissue counterstaining and image segmentation techniques for accurate, quantitative immunohistochemistry.” J Histochem Cytochem 56(10): 873–880.

Zhao, Y., et al. (2018). “TREM2 Is a Receptor for beta-Amyloid that Mediates Microglial Function.” Neuron 97(5): 1023–1031 e1027.

Zhou, S. L., et al. (2019). “TREM2 Variants and Neurodegenerative Diseases: A Systematic Review and Meta-Analysis.” J Alzheimers Dis 68(3): 1171–1184.

Zhou, Y., et al. (2018). “TREM2-Dependent Effects on Microglia in Alzheimer’s Disease.” Front Aging Neurosci 10: 202.

